# Aerosol delivery of immunotherapy and Hesperetin-loaded nanoparticles increases survival in a murine lung cancer model

**DOI:** 10.1101/2024.08.30.609714

**Authors:** Sayeda Yasmin-Karim, Geraud Richards, Amanda Fam, Alina-Marissa Ogurek, Srinivas Sridhar, G. Mike Makrigiorgos

**Affiliations:** Department of Radiation Oncology, Dana-Farber Cancer Institute and Brigham and Women’s Hospital, Harvard Medical School, Boston, Massachusetts, USA; Department of Biochemistry, Northeastern University, Boston, Massachusetts, USA; Department of Biology, Northeastern University, Boston, Massachusetts, USA; CaNCURE program, Northeastern University, Boston, Massachusetts, USA; Department of Physics, Department of Bioengineering, Department of Chemical Engineering, Northeastern University, Boston, Massachusetts, USA

**Author notes:** Correspondence should be addressed to G.M.M. or SYK at Dana Farber Cancer Institute, 450 Brookline Avenue, Boston MA 02115, USA; or.

**Keywords:** Flavonoids, Hesperetin, Nanoparticles, immunotherapy, aerosol drug delivery

## Abstract

**Purpose:** Studies have shown that flavonoids like Hesperetin, an ACE2 receptor agonist with antioxidant and pro-apoptotic activity, can induce apoptosis in cancer cells. ACE2 receptors are abundant in lung cancer cells. Here, we explored the application of Hesperetin bound to PLGA-coated nanoparticles (Hesperetin-nanoparticles, HNPs), and anti-CD40 antibody as an aerosol treatment for lung tumor-bearing mice.

**Methods:** *In-vitro* and in-vivo studies were performed in human A549 (ATCC) and murine LLC1 (ATCC) lung cancer cell lines. Hesperetin Nanoparticles (HNP) of about 60nm diameter were engineered using a nano-formulation microfluidic technique. A syngeneic orthotopic murine model of lung adenoma was generated in wild (+/+) C57/BL6 background mice with luciferase-positive cell line LLC1 cells. Lung tumor-bearing mice were treated via aerosol inhalation with HNP, anti-CD40 antibody, or both. Survival was used to analyze the efficacy of aerosol treatment. Cohorts were also analyzed for body condition score, weight, and liver and kidney function.

**Results:** Analysis of an orthotopic murine lung cancer model demonstrates a differential uptake of the HNP and anti-CD40 by cancer cells relative to normal cells. A higher survival rate, relative to untreated controls, was observed when aerosol treatment with HNP was added to treatment via anti-CD40 (p<0.001), as compared to CD40 alone (p<0.01). Moreover, 2 out of 9 tumor-bearing mice survived long term, and their tumors diminished. These 2 mice were shown to be refractory to subsequent development of subcutaneous tumors, indicating systemic resilience to tumor formation.

**Conclusion:** We successfully established increased therapeutic efficacy of anti-CD40 and HNP in an orthotopic murine lung cancer model using inhalation-based administration. Our findings open the possibility of improved lung cancer treatment using flavonoids and immuno-adjuvants.

## INTRODUCTION

Lung cancer represents the second most common type of cancer worldwide and the leading cause of cancer death [1], accounting for about 1 in 5 of all cancer deaths [2]. Each year, more people die of lung cancer than of colon, breast, and prostate cancers combined [3]. Overall, the chance that a man will develop lung cancer in his lifetime is about 1 in 16; for a woman, the risk is about 1 in 17 [4]. Radiation is a standard modality for lung cancer treatment [5]. Radiation induces DNA damage and apoptosis, causing cancer cell fragmentation that exposes cancer-associated antigens to the tumor microenvironment. These can be recognized by antigen-presenting cells (APC) to induce the antineoplastic effect by activating cytotoxic T cells [6]. In the last few years, there has been a growing interest in cancer immunotherapy due to its promising results in achieving significant and durable treatment responses with manageable toxicity in several cancers including lung cancer [7]. Our prior studies show that adding immunoadjuvants like anti-CD40 antibody along with radiation can enhance anti-tumor (M1) APC cell activation [8]. While beneficial, the use of radiation is also limited by toxicity to normal tissues. For example, when centrally located lung tumors are irradiated, the total radiation dose that can be safely administered is restricted to control radiation toxicity in nearby organs at risk [5].

Flavonoids like Hesperetin, an ACE2 receptor agonist [9, 10], can also induce apoptosis in cancer cells [11], whereas ACE2 receptors are overexpressed in lung cancer [12, 13]. Hesperetin induces cancer cell apoptosis via complex mechanisms invovling targeting of multiple biomolecules and the associated signaling pathways, such as ASK1/JNK, p38/MAPK, Notch1, ROS, Bcl□2 family members, and death receptors [14-18]. Accordingly, Hsperetin treatment can potentially be used with immunotherapy, either as an alternative to radiation, or in combination with a moderated amount of radiation.

This project aims to evaluate an in-vivo inhalation-based drug delivery approach combining nanotechnology with immunotherapy for lung cancer treatment and reduced organ toxicity. We engineered nanoparticles loaded with Hesperetin (HNP) and employed a setup that delivers HNP along with free immuno-adjuvant anti-CD40 antibody as an aerosol preparation for inhalation by mice. Key advantages of using this technique include (a) Increased bioavailability of Hesperetin, delivered in the form of biodegradable HNP. While administering free flavonoids can lead to suboptimal bioavailability, nanoparticle-based delivery may partly overcome this limitation [19] and enhance the binding of Hesperetin to lung cancer cell receptors. (b) Providing a noninvasive alternative to intravenous (IV) or intramuscular (IM) injections [20]. And (c) in-situ delivery of HNP and anti-CD40 into lung tumors. Due to the large surface area and high vascularization, the lungs are an attractive delivery route for treating respiratory and systemic diseases [21, 22], bypassing degradation in the gastrointestinal tract and first-pass metabolism in the liver [22]. Standard intravenous delivery results in less than 1-5% of the drug reaching the tumor [23], thus the proposed local delivery can be anticipated to boost drug delivery into tumors and improve tumor/non-tumor ratios.

## MATERIALS AND METHODS

### Cell line and Molecules

Luciferase gene labeled LL/2/Luc-2, Lewis lung carcinoma cell line was purchased from ATCC and was cultured in Dulbecco’s Modified Eagle’s Medium (DMEM) (GIBCO) with 10% FBS (Sigma) and 1% penicillin/streptomycin (Invitrogen). Red cherry Fluorescent tagged Anti-CD40 was purchased from Abcam.

### Microfluidics-based method to synthesize PEGylated nanoparticles

Hesperetin (C_16_H_14_O_6_) and polymers were purchased in powder format from Sigma Aldrich and kept at 4-8^0^C. We used a microfluidics-based synthesis protocol to generate Hesperetin-loaded nanoparticles (HNP). A syringe pump (‘Harvard Apparatus’), microfluidics chips, and tubing (Sigma) were used to synthesize nanoparticles of specific sizes, encapsulating the hydrophobic hesperetin within biodegradable PEGylated polymers following the company’s instructions. Acetonitrile and dimethyl sulfoxide (DMSO) were used as hydrophobic solvents to dissolve the hydrophobic flavonoid Hesperetin and polymers. Two polymers were used to increase or decrease the drug-releasing time, PLGLA-L and PLGLA-H, Sigma. Flow rates were adjusted to synthesize different-sized nanoparticles. Dialysis was performed with 2 liters of distilled water to discard the free drug and polymer. The protocol was optimized to generate nanoparticles with varying the polymers PLGA50-L and PEGPLA-L and flow rates. We synthesized different sizes of NPs using different flow rates (100 - 470 μl/min) in the presence or absence of Hesperetin in various concentrations (400-800 ng/ml). A plate reader was used to measure the HP concentration of the prepared HNPs. Prepared nanoparticles were imaged with Transmission Electron Microscopy, TEM, at the Northeastern University TEM microscopy facility. Samples were collected within 24 hours after the nanoparticle preparation. All prepared nanoparticles were kept at 4-8^0^C.

### Computed Tomography (CT) Imaging

CT imaging was performed using a small animal radiation research platform (SARRP, Xtrahl, Inc.). Mice were anesthetized with isoflurane vapor, and CT images were taken at 65 kVP energy.

### Pathology and Histology

Ex-vivo tissues were collected and fixed with 10% formalin for 24 hours. Tissues were processed for paraffin embedding by iHisto Co. (Massachusetts). For gross pathology to identify lung tumor prognosis. For Hematoxylin and Eosin (H&E) Staining, Paraffin-embedded tissues were sliced into four µm-thick sections with a microtome, air-dried, fixed with acetone, and stained following standard protocol. Sections were stained with hematoxylin and eosin (H&E) to observe lung metastasis, and whole slide scanning (40x) was performed on an EasyScan infinity (Motic). High-magnification images were collected using Case Viewer software.

### Mouse and Tumor Models

C57BL/6NTac background 8-12 weeks-old wild (W^+/+^) male and female mice were purchased from Taconic mice. The animals were contained in groups of five in standard cages with free access to food and water and a 12□h light/dark cycle. All mice were adjusted to the animal facility for at least 1□week before experimentation. All possible parameters that may cause social stress, like group size, among the experimental animals were carefully checked and evaded. Animals were observed three times a week after cell implantation for any physical abnormalities. To better mimic the lung cancer scenario, orthotopic lung tumors were generated by implanting cancer cells in mouse lung using an endotracheal intubation kit (Kent Scientific Kit). C57BL/6 background mice were implanted with the matched background LL/2-Luc2 (10 ×10^5^ cells/tumor) luciferase-positive cell line (ATCC). The luciferase-expressing mouse cells are ideal for in vivo bioluminescence imaging to monitor NSCLC tumor development, in vivo.

For subcutaneous (SQ) tumor models, LL/2-Luc2 (5 ×10^4^ cells/tumor) suspended in PBS were injected into one of the mice’s flanks. In all cases, an insulin syringe with a 22-gauge needle was used for subcutaneous tumor cell injection. All mice were maintained following an IACUC-approved protocol and complied with the guidelines and regulations set by the Institutional Animal Care and Use Committee (IACUC).

### Aerosol drug delivery

We developed a mice aerosol drug delivery system using a hospital-grade Schuco air compressor, a mouse drug delivery mixing chamber, and a mouse pie cage for a nebulizing drug delivery system (Braintree Scientific, Inc). The mouse pie case is designed for aerosol drug delivery of up to 10 mice at a time. Aerosol enters the pie via tubing attached to the central core. The core has a sealable opening apparatus on the top for mouse placement and a solid on the bottom. Aerosol exits each pie section through one 3 mm hole in the center of the outer edge. According to the company, the Nebulizer Compressor uses compressor pressure of 35-45 psig (pound per square inch gauge), with an operating flow rate of 7-11 Ipm (inch per minute), releasing therapeutic particle size - 0.5 µm-5µm using AC operation -of 120V 60 Hz. Using this aerosol drug delivery system, we successfully treated lung tumor-bearing mice with HNP of 500ng/ml (250ng/Kg) solution using PLGA50-L polymer and/or anti-CD40 (40µg/ml) diluted to 5 ml with sterile ddH_2_O. We used a lower dose (less than the average (< [(400+800)/2]) dose of treated SQ tumors to reduce the drug-related complications). We used same-sized drug-free PLGA50-L polymer NPs for control mice with a similar dilution. Aerosol delivery was performed for 30 minutes. For all cohorts, body weight and body condition scores were measured. Body conditions were scores following the DFCI IACUC guidelines (one being the lowest and four being the highest score based on body fat condition).

### Statistical Analysis

Survival data was plotted, and statistical analyses were performed using GraphPad prism v7.0. A log-rank test was employed to determine the p-value for the Kaplan–Meier curves. Statistical analyses for tumor volume were achieved using two-way ANOVA: two factors with replication tool and standard Student’s two-tailed t-test. P-values <0.05 were considered significant. *P□<□0.05, **P□<□0.01, ***P□<□0.001, and ****P□<□0.0001 were considered as statistically significant at 95% confidence interval.

## RESULTS

We first analyzed the effect of Hesperetin (**Figure 1A**), on the proliferation of human A549 lung cancer cell line, using a clonogenic survival assay, showing a significant decrease in cell survival with 5 (p<0.01), 15 (p<0.001), and 25 (p<0.0001) µg/ml of Hesperetin treatment for 24 hours, compared to the control. The cell survival percentage decreased in a dose-dependent manner, with an IC50 value of approximately 10 μg/ml concentration of Hesperetin (**Figure 1B**). Next, given the evidence of the anti-cancer cell activity of Hesperetin towards the human lung cancer cell line, using a microfluidic technology (**Supplementary figure 1A**), we generated Hesperetin-loaded Nanoparticles (HNPs) of two different sizes, S and L (**Figure 2A and Supplementary figure 2B**). Attaching Hesperetin to polymer-based nanoparticles was anticipated to improve the bioavailability of the flavonoid in tissues via slow and sustained in-situ delivery of the drug payload [24]. Size of HNP varied depending on polymer used (**Figure 2B**). To analyze the HNPs’ effect on lung cancer treatment, we used an *in-vivo* murine subcutaneous (SQ) lung cancer model by generating tumors on immunocompetent C57BL/6 mice with matched background mouse lung cancer cell line (LL/2-Luc2). Here, we used two different-sized HNP using two different polymers, PLGA-50L (Sigma) and PEGPLA-L for the larger-sized [**L**] (average diameter ∼60 nm, range = 37-105 nm) and smaller-sized [**S**] (average diameter ∼40 nm, range = 31-60 nm) of HNPs, respectively. Palpable-sized SQ tumors were injected directly with two different concentrations of HNPs, 400 ng/ml (200 ng/kg) and 800 ng/ml (400 ng/kg) of HNP solutions for each size (S and L). Total treatment was given three times, three days apart (**Figure 2C)**. Two different sizes of HNPs (S with PEGPLA-L and L with PLGA-50L polymers) were used for each concentration of HNPs. The SQ tumor volume data show that HNP can suppress the growth of lung tumors in mice in a dose-dependent manner (**Figure 2D**).

**Figure 1.**
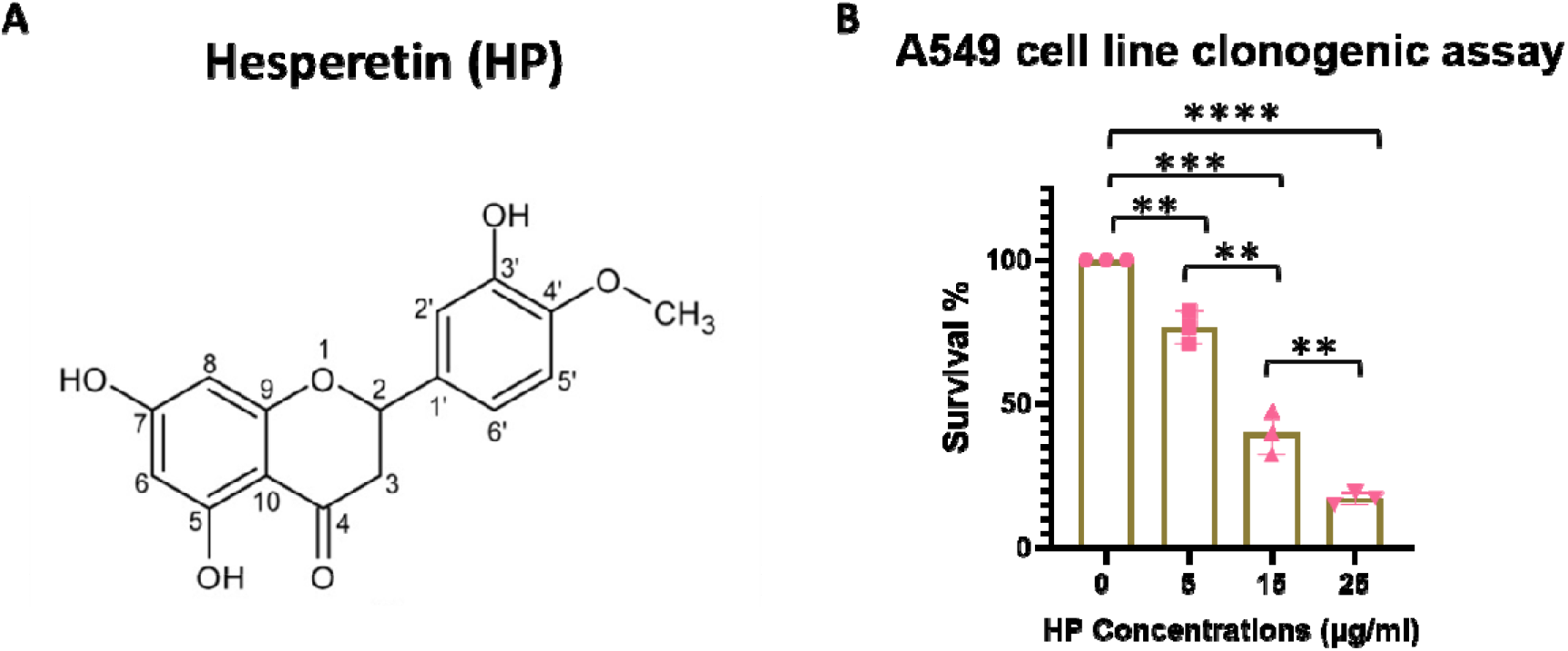
Anti-tumorigenic effect of Hesperetin in human lung cancer cell line. A. Biochemical structure of Hesperetin. B. Bargraph showing clonogenic survival assay (n=3) in human lung cancer cell line, A549, treated with Hesperetin doses from 5 to 25 µg/ml for 24 hrs. Data represent the mean +/-standard deviation (SD) of three experiments. ** p< 0.01, *** p<0.001, **** p<0.0001 at 95% confidence interval.

**Figure 2.**
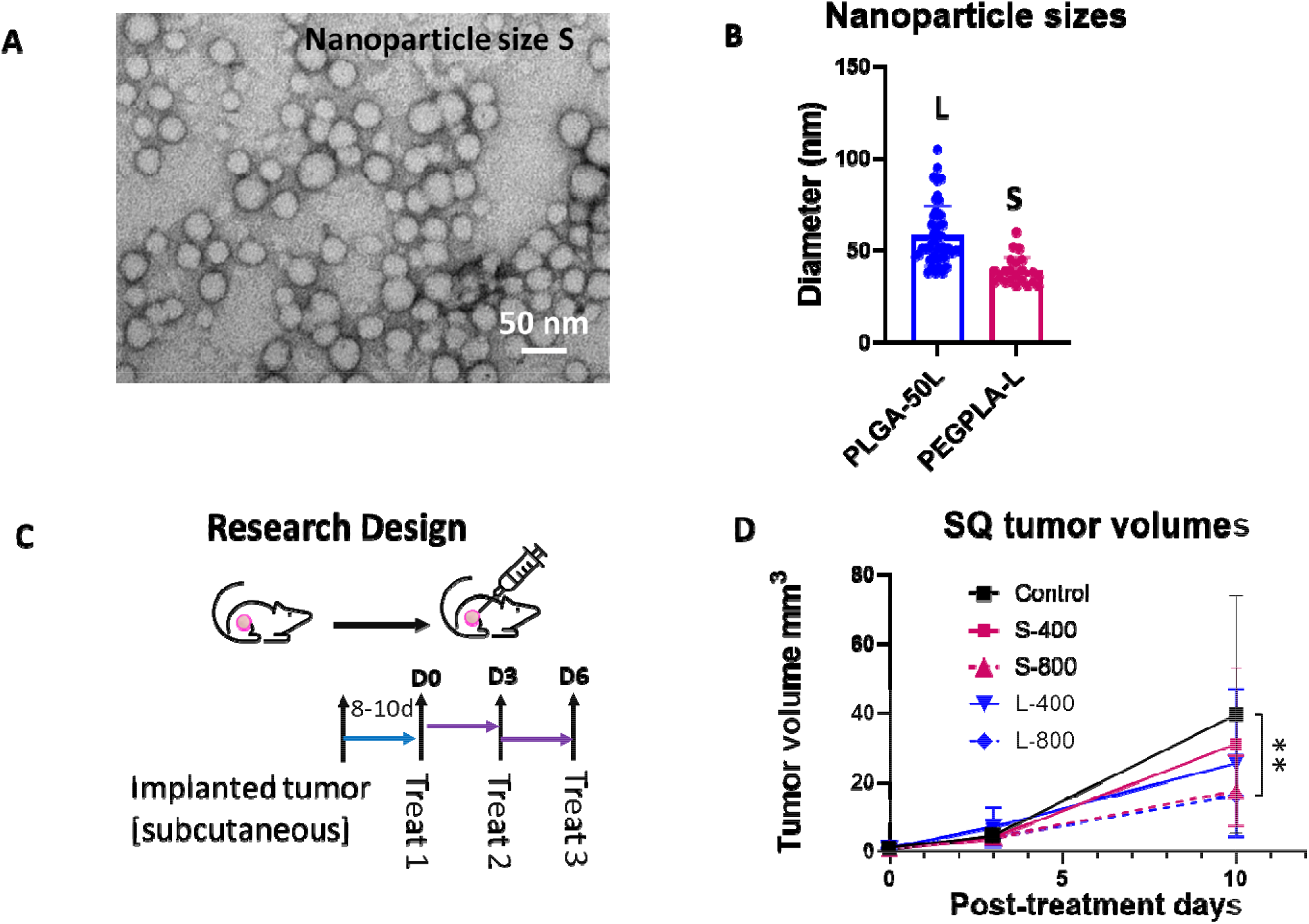
Preparation of Hesperetin-loaded nanoparticles (HNP) and efficacy in a subcutaneous (SQ) mouse tumor model. A. A Transmission Electron Microscopy (TEM) image of HNP (representative of size S) and B. Bar graph of the different-sized (S and L representing 31-60 nm and 37-105 nm in diameter, respectively) HNPs used in the efficacy study in the SQ lung tumor model. C. Study design and workflow. D. Tumor growth curve for different sizes and concentrations of HNP treatment in an SQ mouse model for mouse lung cancer showing up to 10 days while control mice were still alive. A single SQ cancer was implanted in one of the flanks of C57BL/6 background immunocompetent mice with matched background mouse LLC1 cell line, and all cohorts (n=5) were treated when the tumor reached palpable size. Control mice were treated with the same vehicle volume (100 µl). Data represents the mean +/-SD. * p< 0.5 at 95% confidence interval.

To further evaluate the in-vivo action of HNP in murine lung cancer, we generated a lung orthotopic mouse cancer model and followed the growth with BLI imaging (**Figure 3A**). We used intubation to implant the luciferase reporter-labeled cells LL/2-Luc2LLC-1 on the same C57BL/6 mouse background lung cancer cell line, to generate orthotopic lung cancer, one tumor per mouse (**Supplementary Figure 2A and B**).

**Figure 3.**
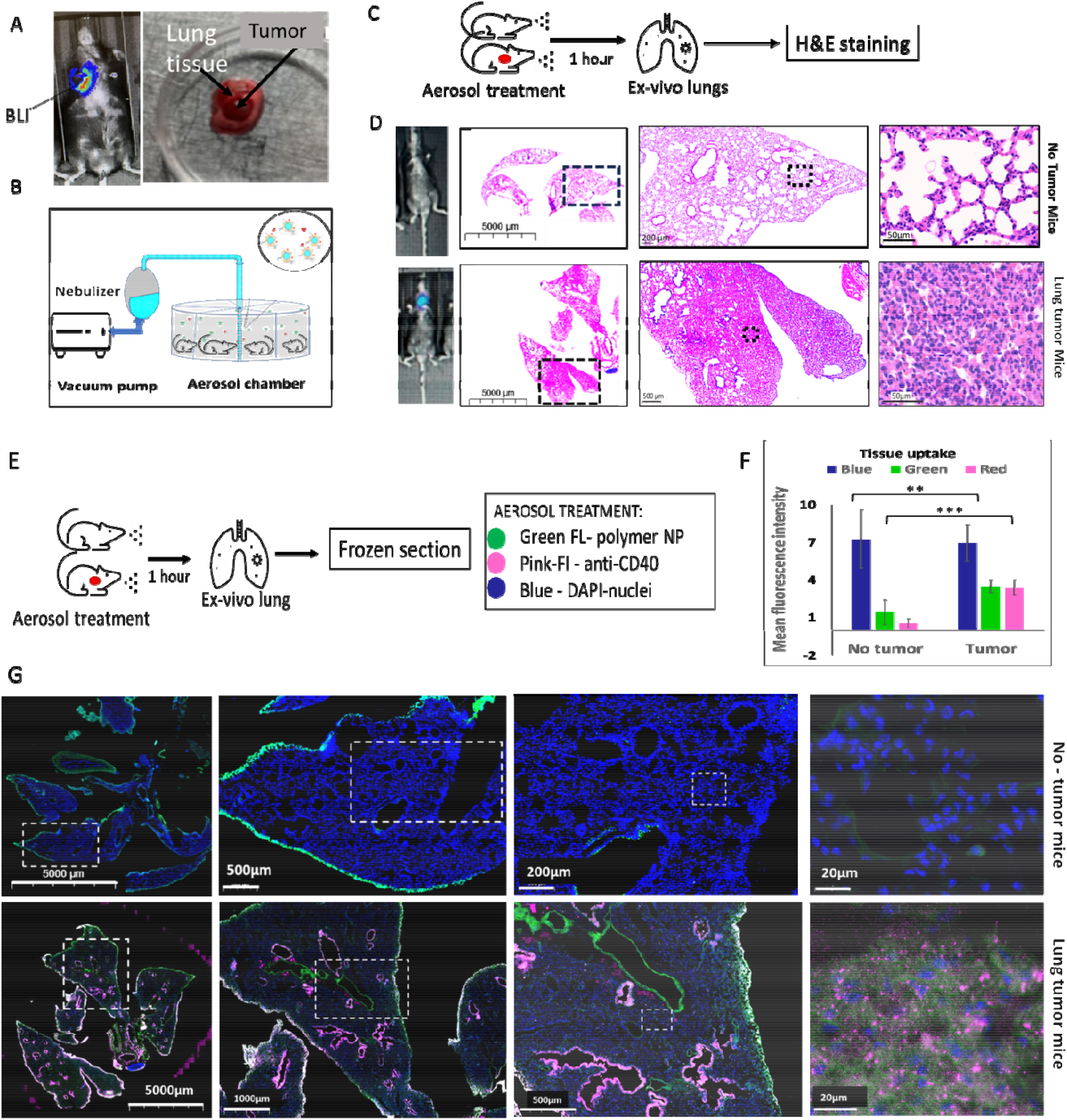
Establishment of Aerosol delivery of HNP and anti-CD40 in Lung Cancer Orthotopic animal model. A. BLI image (left) of a mouse showing the development of lung tumor by the implantation of mouse lung cancer cells (LLC1) under the guidance of endotracheal intubation tubing. (Right) ex-vivo picture of tumor grown in lung tissue. B. Schematic diagram of the aerosol drug delivery using a nebulizer vacuum pump connected to a drug mixture chamber delivering to an aerosol drug delivery chamber. C. Treatment design for the aerosol drug delivery of HNP and free antiCD40 in mice followed by hematoxylin and eosin (H&E) staining in ex-vivo lung tissue in naïve and tumor bearing mice. D. Bioluminescence (BLI) images of the mice and H&E staining images of the ex-vivo lung tissue mice under different magnifications show no tumor (upper) and lung tumor-bearing mice (bottom). E. Research design for aerosol drug delivery of FITC labeled polymer NP (green, wavelength ∼520 nm) and red mcherry dye fluorescently tagged antiCD40, (red, wavelength ∼600 nm). Cells are represented by DAPI (blue wavelength ∼460 nm) staining of the nuclei. F. and G. Bar graph and fluorescence images, respectively, of ex-vivo lung tissue depicting cell nuclei stained by DAPI (blue), FITC labeled Polymer NP (green) uptake and red mcherry-tagged Anti-CD40 antibody uptake (red-pink). Diffuse co-localization of fluorescent nanoparticles and anti-CD40 was observed in tumor-bearing mice (upper panel row) but not in the tumor-free mice (lower panel row), in the tumor microenvironment.

We then employed an aerosol drug delivery system using a nebulizer (**Figure 3B and Supplementary Figure 3**) to establish an inhalation drug delivery system for lung cancer treatment. Using this aerosol inhalation system, we initially employed green fluorescence polymer nanoparticles (NPs) to confirm the uptake of NPs by lung tissue when using the aerosol drug delivery system. In addition, we used red-cherry fluorescence tagged Anti-CD40 for inhalation treatment along with green fluorescence polymer NPs. To observe the uptake in tumor-inoculated lungs, we used lung orthotopic tumor-inoculated mice and healthy mice. Tumor formation was initially observed via BLI imaging and later confirmed by ex-vivo H&E image of the resected lungs (**Figure 3A-D**). Two weeks after tumor inoculation, both groups of mice were given aerosol delivery of green fluorescence-tagged polymer NP and red fluorescence-tagged Anti-CD40 for 30 minutes simultaneously. Ex-vivo resection of the lung’s frozen section (**Figure 3E**) within an hour following the treatment demonstrated preferential uptake of both HNP and anti-CD40 (bar graph **Figure 3F)** by lung tumor tissues (images **Figure 3G**, lower panels) as compared to healthy lung tissues (**Figure 3G**, upper panels).

Following confirmation of nanoparticle and anti-CD40 uptake in lung tumor cells, we conducted experiments to observe the therapeutic effect of HNP along with free Anti-CD40 following aerosol drug delivery (**Figure 4A**). The data demonstrate an increased survival rate and duration in the case of tumor-bearing mice treated with free anti-CD40 inhalation compared to the placebo group (**Figure 4B**). Further survival enhancement was observed when HNP was added along with anti-CD40 during aerosol therapy, as shown by the Kaplan-Meyer survival graph (**Figure 4B**). Representative BLI images are shown in **Figure 4C**. Ex-vivo H&E staining of the lung was performed in a fraction of mice, showing tumor regression in representative images (**Figure 4D**).

**Figure 4.**
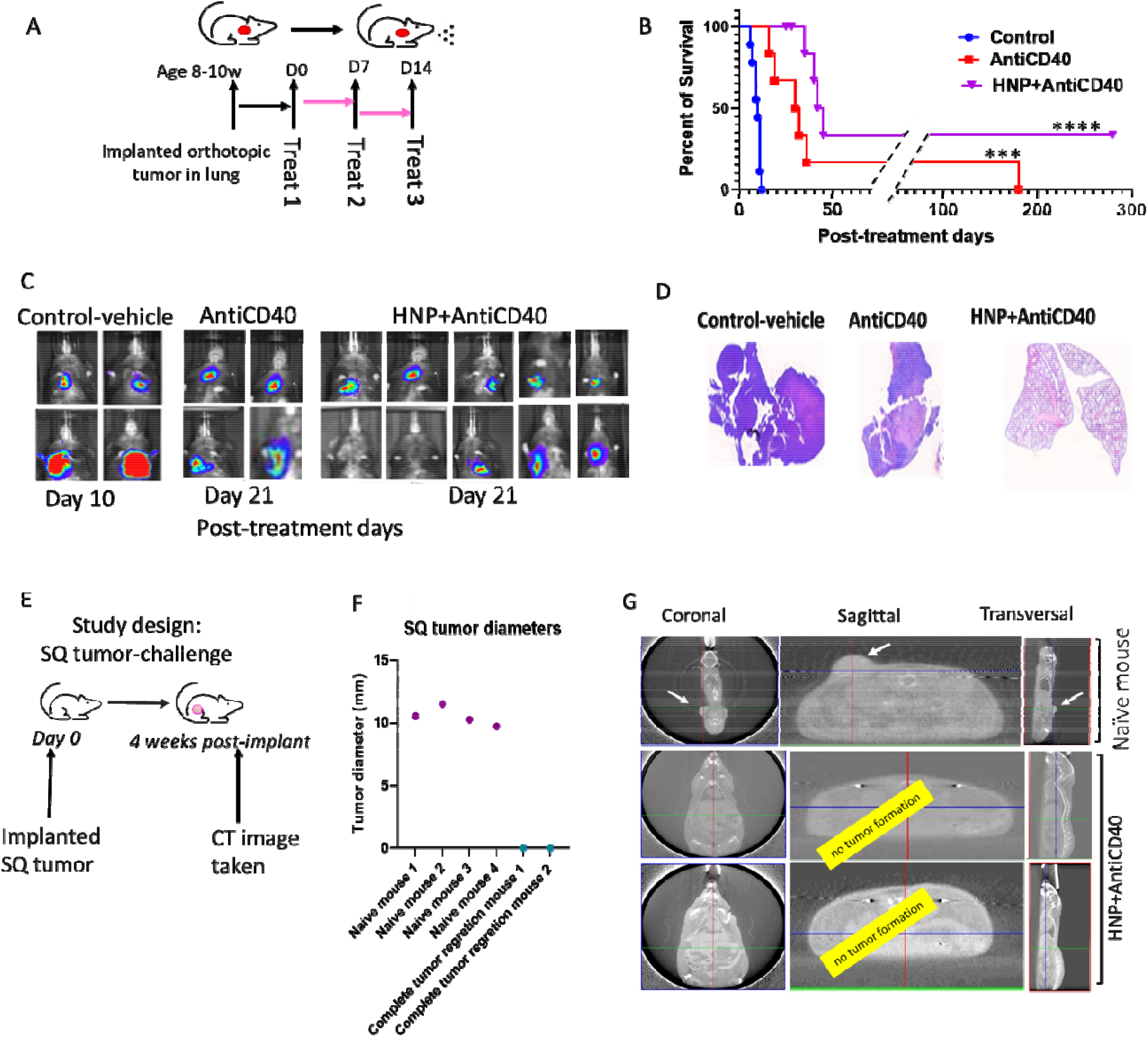
Efficacy of HNP and anti-CD40 in Lung Cancer Orthotopic animal model with Aerosol drug delivery. A. A schematic diagram of the aerosol drug delivery in orthotopic lung cancer bearing mice. B. Kaplan Meier survival graph showing the survival percent and duration of different cohorts of treated mice with aerosol delivery of AntiCD40 (n=6) and HNP+AntiCD40 (n=9) with control (n=8). C. Representative BLI images of the different cohorts from the treated mice and untreated controls; D. Representative H&E images from the ex-vivo lungs confirming the corresponding treatment outcome. E. Study design. F. Graph showing the outcome of rechallenging the recovered mice with an SQ implant of LLC1 tumor cells along with a cohort of naive mice as controls (n=4). G. CT images taken four weeks post-injection, confirming the tumor growth outcome in naïve (n=4) and HNP+AntiCD40 treated mice that had complete tumor regression (n=2). For statistical significance at a 95% confidence interval, *** p< 0.001, **** p< 0.0001.

Two of the nine treated mice with combination therapy demonstrate complete tumor regression with a survival duration of more than 300 days post-treatment. A subsequent re-challenge of these two long-surviving mice was performed with subcutaneous tumor cell implantation using the same LLC1 cell line (**Figure 4E**) and demonstrated no tumor growth in the surviving combination-treated group. In contrast, all 4 -naive mice with subcutaneously implanted LL/2-Luc2 cells developed SQ tumors as shown in manually measured tumor diameter (**Figure 4F**) and by CT images (**Figure 4G**). The data are consistent with a ‘vaccine effect’ development in the two long-surviving mice from the combination-treatment group.

The toxicity study demonstrates no change in body score and body weight in this treatment cohort (**Figures 5A and B**), and no significant deviation in the liver and kidney function for aerosol HNP treatment after three hours (**Figures 5C**) and three days post-treatment (**Figure D**) when compared to mouse upper and lower range of normal serum levels (red arrows).

**Figure 5.**
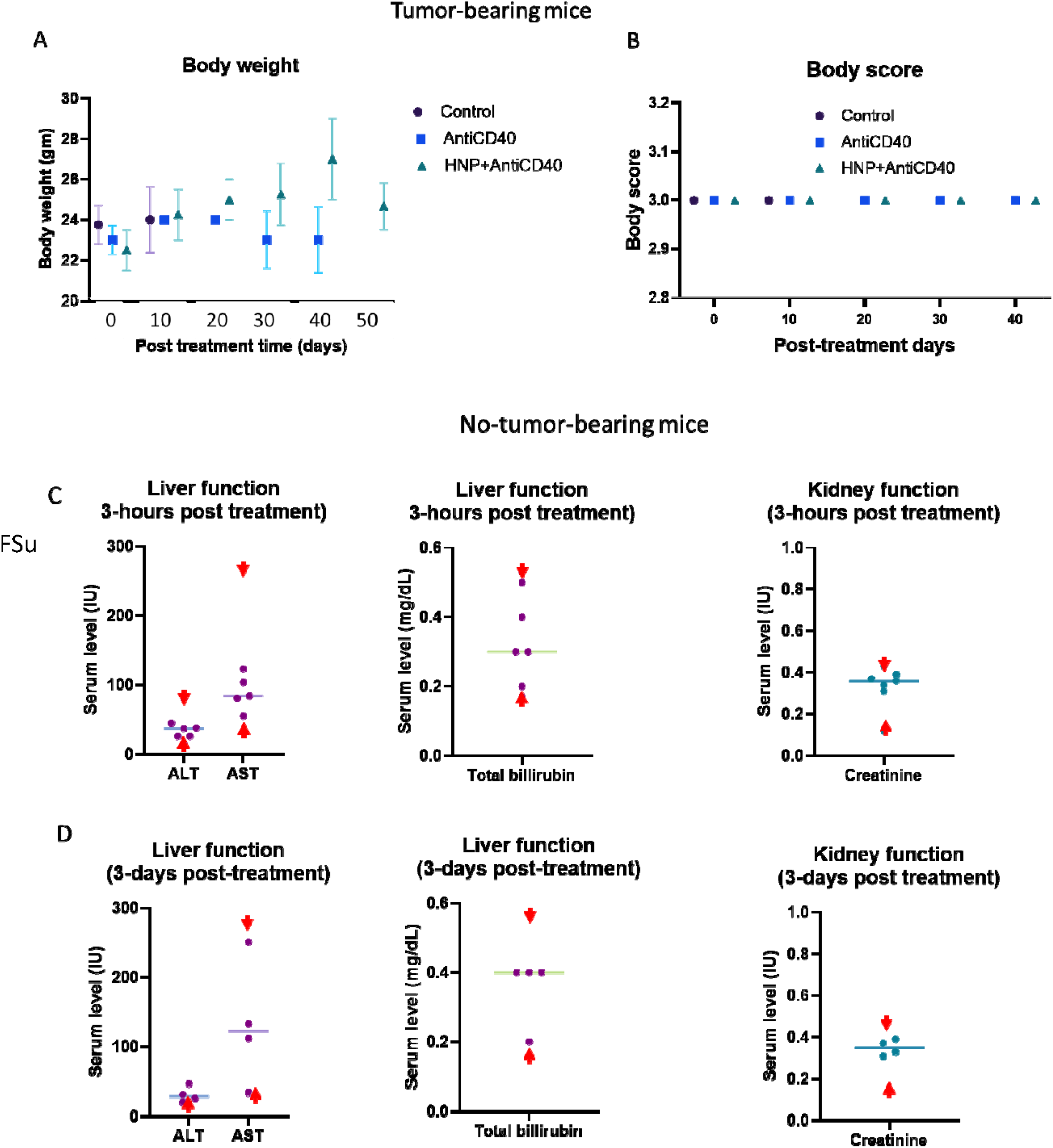
Toxicity study. A. Body weight and B. body score were measured during the treatment period of the tumor-bearing mice for all cohorts of the aerosol-treated mice along with the controls. Each data represents the average outcome of the cohort for that time. C. Three hours (n=5) and D. three days (n=4) post-treated liver and kidney function clinical reports of the combination-treated (HNP+AntiCD40) mice with aerosol-treated doses in wild naïve (non-tumor bearing) mice. Ex-vivo serum was collected for clinical laboratory reports for analysis of serum alkaline transaminase (ALT, normal range 15-80.1 U/L), serum aspartate transaminase (AST, normal range 33.95-268.47 U/L), serum bilirubin (normal range 0.17-0.53 mg/dL), and serum creatinine (normal range 0.12-0.43 mg/dL). Each data represents an individual mouse. Red font arrows (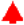 and 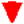) indicate the mouse’s upper and lower range of normal serum levels.

## DISCUSSION

Our ability to manipulate and manufacture materials on the nanoscale level allowed us to develop nanotechnology – based approaches combined with radiation therapy, chemotherapy or immunotherapy. In earlier work, we proposed the replacement of fiducials used in radiation therapy for guiding radiation beams to the tumors with gold nanoparticles loaded with radio-sensitizers and encapsulated in slow-release polymers, to enable simultaneous beam localization and biological sensitization [25, 26]. This approach can be used both for uniform, externally applied radiation therapy as well as for non-uniform radiation therapy delivered via brachytherapy or internal emitter radiation therapy [26-30]. In the present work, we employed nanotechnology-based drug inhalation strategies for improving therapeutic efficacy and mitigating adverse side effects in lung cancer harboring mice. Polymeric nanoparticles remain stable during nebulization and induce minimal toxicity, while having tunable degradation rates in-vivo [22]. Our experimental protocol proved effective in introducing hesperetin onto nanoparticles, inhaling the nanoparticles, and eventually releasing hesperetin in lung cancer tissues where, along with free immunoadjuvant anti-CD40, reduced or eliminated growth of implanted lung tumors.

Lung cancer evades immune response through multiple mechanisms. For instance, lung cancer cells may undergo immunoediting, in which precancerous cells slowly undergo selective adaptation to become ‘invisible’ to immune surveillance [23]. Previous studies [24, 25], including work from our group [8] demonstrated that immunoadjuvants like anti-CD40 induce antitumor effects in different cancer models, including the lung, while apoptosis induced by radiation initiates further enhancement of immune cell activation [6, 7]. Here we used hesperetin as a pro-apoptotic agent [12], instead of radiation. We show that the addition of hesperetin-loaded nanoparticles to the immunoadjuvant anti-CD40 and delivery in aerosol format is effective in treating orthotopic lung cancer in mice, even without the inclusion of radiation. In addition, there is evidence that a fraction of the mice (2/9) obtained long-term immunity that prevented subcutaneous tumor formation upon re-challenging with the same tumor cell line. The data are consistent with an abscopal, cancer-vaccine effect [26], upon which endogenous T cells are primed against the cell line tumor antigens following the applied aerosol treatment. While the abscopal effect hypothesis must be validated with additional studies, almost all treated mice exhibited an increase in survival following a relatively simple inhalation treatment that was administered non-invasively, and without advanced equipment. This treatment approach may potentially allow lung cancer treatment that is readily available in under-resourced environments that have reduced access to radiation and without the expense and hazards associated with radiation treatment. Alternatively, the treatment could be offered in conjunction with fewer radiation courses and a reduced total radiation dosage. We anticipate that nanoparticle-based aerosol drug delivery locally for lung cancer, potentially in conjunction with locally administered radiation, may enhance the overall efficacy of treatment while drastically reducing systemic treatment complications.

## Acknowledgements

This work was funded by NCI grant 2R25 CA174650 (CaNCURE program, Northeastern University), the JCRT Foundation, and Nanocan Inc.

## Authors approvals

all authors have seen and approved the manuscript. The manuscript has not been accepted for publication or published in a different journal.

## FIGURE LEGENDS

**Supplementary Figure 1.** A. Engineering of nanoparticles: Apparatus used for microfluidic generation of Hesperetin-loaded nanoparticles. B. Transmission Electron Microscopy (TEM) image of HNP demonstrate their size distribution, L.

**Supplementary Figure 2**. Development of the orthotopic lung tumor model by endotracheal intubation using an endotracheal tumor implant tubing kit for mice. A. Light-guided tubing system for the tumor cell implantation through endotracheal tube to the lung. B. The position of the mice during tumor implantation. Mice were kept in this vertical position for a few minutes following implantation, to help migrate cancer cells toward lung tissue, via gravity.

**Supplementary Figure 3**. Establishment of aerosol delivery of HNP and Anti-CD40 in lung cancer orthotopic animal model. The picture shows the aerosol drug delivery in mice located in a multi-pocket chamber for the treatment of up to 10 mice at a time connected to a drug-mixing chamber with a nebulizer system for aerosol inhalation.

