## Supplementary figures and images for "Aerosol delivery of immunotherapy and Hesperetin-loaded nanoparticles increases survival in a murine lung cancer model"

### supplemental figures 1-3

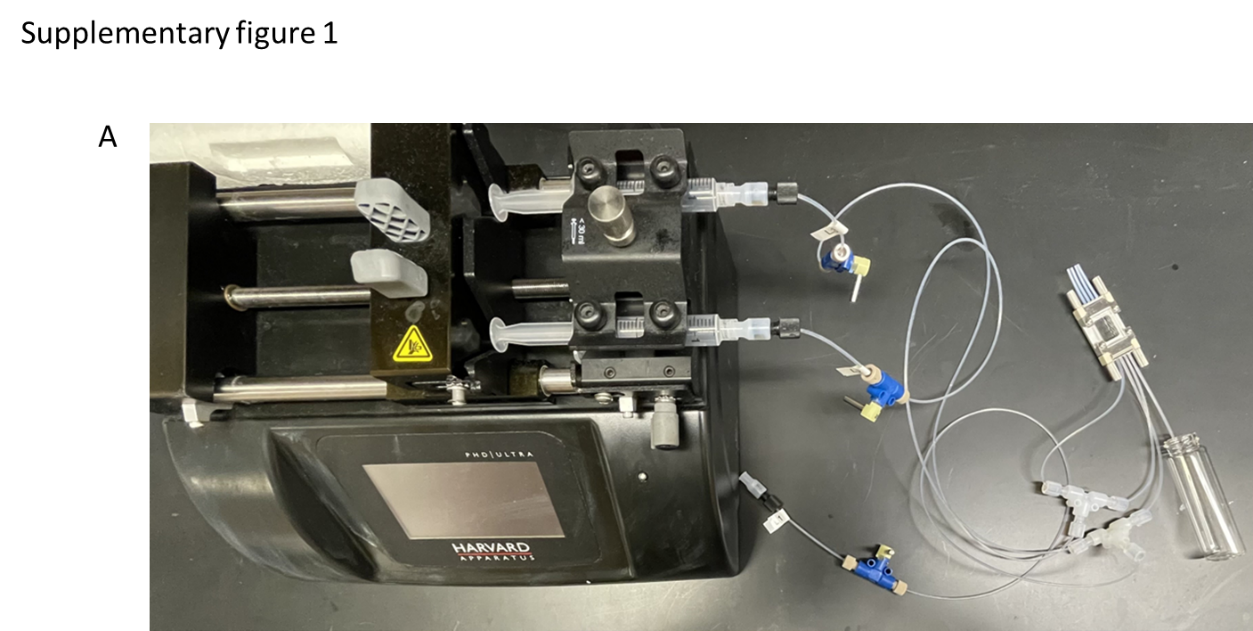

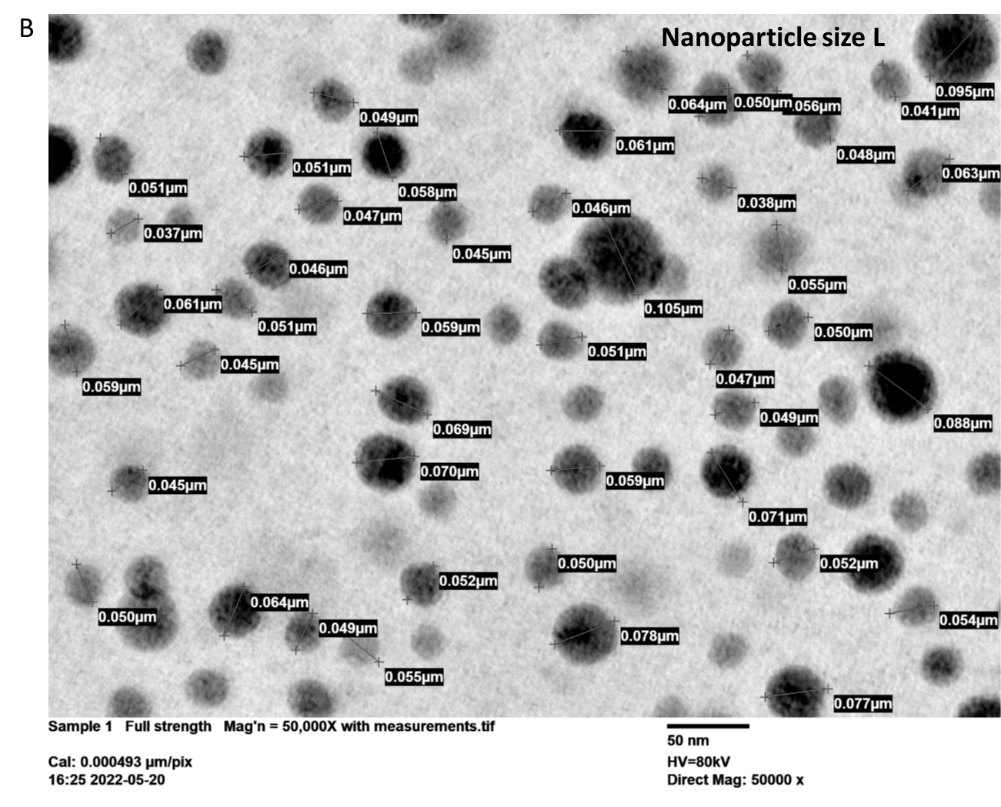

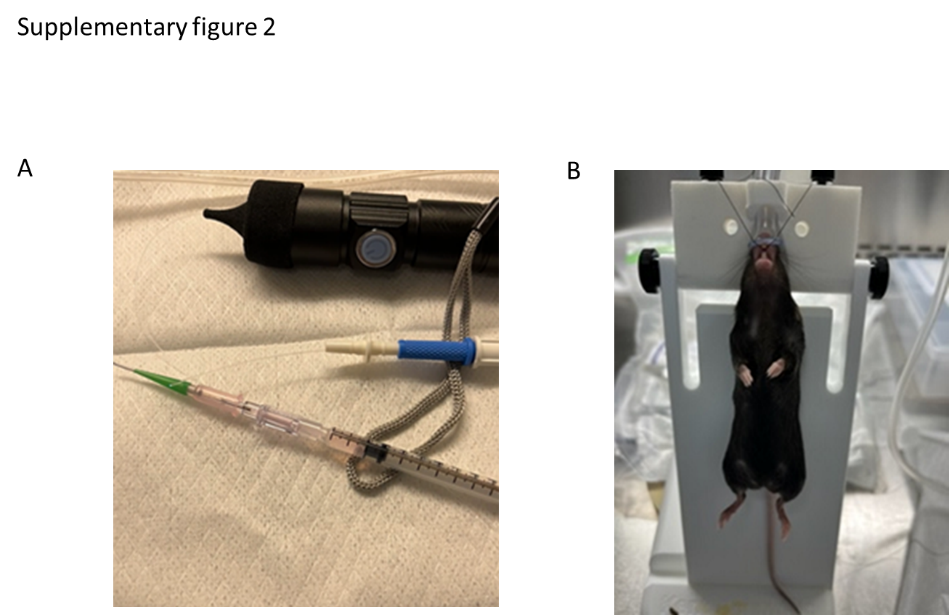

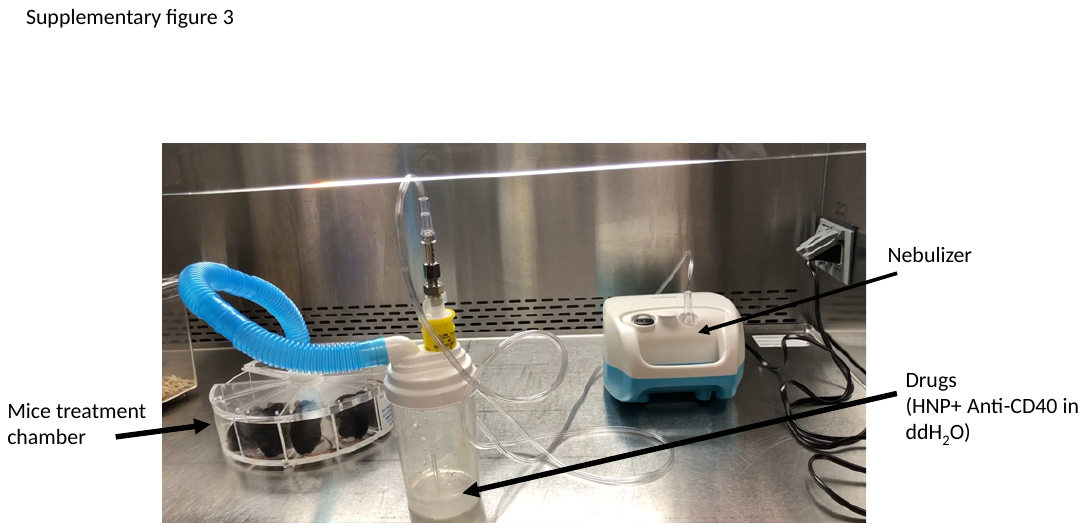
